# Functional Conservation Amid Extreme Divergence in Freshwater SAR11 Hypervariable Region 2

**DOI:** 10.64898/2026.11.09.765219

**Authors:** Shaanaav Daniel, Adrian-Stefan Andrei, Yohei Nishikawa, Haruko Takeyama, Hiroyuki Ogata, Yusuke Okazaki

## Abstract

The ‘Ca. Pelagibacterales’ order (SAR11) is a ubiquitous bacterioplankton in aquatic ecosystems. The genomes of SAR11 bacteria contain hypervariable region 2 (HVR2), the largest HVR with extreme diversity. Previous studies collectively indicated the enrichment of cell-surface modification genes on HVR2, which have been hypothesized to facilitate phage evasion. However, characterisation of HVR2 has primarily targeted marine clades and been limited for freshwater SAR11 (aka. *Fontibacterium*). Here, we employed single-cell genomics and recovered 176 *Fontibacterium* HVR2 sequences from Lake Biwa (Japan) and Lake Zurich (Switzerland), across multiple sampling years for Lake Biwa. Comparative sequence analyses revealed that freshwater HVR2 showed high intra-population diversity and appears to evolve independently of the rest of the genome, mirroring patterns observed in marine counterparts. Nevertheless, highly conserved HVR2 sequence pairs were observed for distinct *Fontibacterium* species from the two intercontinental habitats, suggesting horizontal transfers of HVR2 with conserved functional configurations across divergent lineages and large spatial scales. Further comparison with 145 marine HVR2 sequences revealed HVR2 sequences displaying synteny between *Fontibacterium* and marine SAR11, and 209 gene families conserved across habitats, including three near-universally conserved families. All examined freshwater HVR2s contained initiator methionine tRNA genes concentrated near the boundaries of the 5S region, suggesting a tDNA-anchored integration mechanism. These results suggest the functional equivalence of HVR2 across marine and freshwater SAR11 clades, and the existence of a shared pool of HVR2 sequences that evolve reticulately through mutual genetic exchange, independent of the vertical inheritance of the rest of the genome.

## INTRODUCTION

The SAR11 clade (the order Pelagibacterales) is the most abundant heterotrophic bacterial group in aquatic ecosystems, accounting for over 25% of total bacterioplankton in global surface oceans (Giovannoni, 2017; Morris et al., 2002) and freshwater lakes (Heinrich et al., 2013, Salcher et al., 2011). Due to its extreme abundance, SAR11 is a key driver of the biogeochemical cycles in marine ecosystems. The SAR11 clade is phylogenetically diverse with multiple subclades, each occupying a distinct ecological niche. In freshwater lakes, the subclade IIIb/*Fontibacterium* (formerly “*Candidatus* Fonsibacter”) is the dominant lineage (Fernandes et al., 2025). Marine SAR11 members belong to subclades which are phylogenetically distinct from the genus *Fontibacterium*. Despite their environmental dominance, SAR11 remains difficult to culture with only a few reports of successful isolation (Fernandes et al., 2025, Henson et al., 2018, Salcher et al., 2025), making their ecophysiological characterisation still limited. The known diversity of SAR11 has been mostly characterised from genomes generated by cultivation-independent omics approaches.

One of the major features of the SAR11 genomes is a large number of hypervariable regions (HVRs) (Molina-Pardines et al., 2025), which are genomic regions with elevated variability relative to the conserved core genome. The largest of these is HVR2, a ∼48kb region flanked by the 16S–tRNA– tRNA–23S cassette on one side and the 5S rRNA gene on the other (Wilhelm et al., 2007), a feature that has been conserved in freshwater SAR11 (Fernandes et al., 2025). The rRNA regions potentially act as anchors for homologous recombination (Wilhelm et al., 2007), which could allow HVR2 cassettes to be exchanged between strains. Horizontal gene transfer is also suggested to contribute to HVR2 diversification, as novel cell-surface modification genes have been found to be acquired from distantly related lineages (Zaremba-Niedzwiedzka et al., 2013). Recent works have shown that this diversification operates at the strain level within a single population, with nearly every co-occurring strain carrying a distinct HVR2 (Haro-Moreno et al., 2025; Molina-Pardines et al., 2025). HVR2 is dominated by genes that synthesise cell surface proteins, such as glycosyltransferases and O-antigen polymerases (Grote et al., 2012, Zaremba-Niedzwiedzka et al., 2013, Henson et al., 2018, Chen et al., 2019, Haro-Moreno et al., 2025). These cell-surface modification genes have been suggested to function for defence against phages; by diversifying the displayed polysaccharides on the outer membrane, bacteria can evade receptor recognition and prevent phage adsorption (Rodriguez-Valera et al., 2009). Phage predation pressure has been documented across both habitats. Pelagiphages infect marine SAR11 (Martinez-Hernandez et al., 2022, Zaragoza-Solas et al., 2020, Zhao et al., 2013), while closely related phages infect *Fontibacterium* in freshwater systems globally (Chen et al., 2019, Okazaki et al., 2019). Yet, the extent to which marine and freshwater HVR2s share genes that contribute to antiphage function has not been characterised.

Conventional metagenomic approaches are poorly suited to resolving HVR2 intra-population diversity, because metagenomic assembly capture genomes of co-occurring strains only as a single consensus sequence and therefore cannot resolve microdiversity (Ramos-Barbero et al., 2019). This limitation cannot be overcome even by long-read or deep (high-coverage) sequencing (Okazaki et al., 2022); even when a hypervariable region is recovered by metagenomic assembly, the resulting consensus sequence does not reveal the diversity within that region. Single-cell genomics circumvents this issue by amplifying and sequencing the individual genomes of cells, producing high-resolution single amplified genomes (SAGs). While *Fontibacterium* is difficult to cultivate (Fernandes et al., 2025), a SAG-based approach was considered effective for high-resolution characterisation of HVR2 and the surrounding genome. To date, marine HVR2 data has expanded to over 140 sequences through a combination of metagenomics and single-cell approaches (Haro-Moreno et al., 2025), while freshwater HVR2s have generally been examined at a significantly smaller scale (Fernandes et al., 2025;Okazaki et al., 2025; Zaremba-Niedzwiedzka et al., 2013; Henson et al., 2018; Chen et al., 2019). The shortage of freshwater HVR2 data has left their diversity and evolutionary dynamics as well as the interplay between freshwater and marine HVR2 unclear.

Furthermore, freshwater systems offer a complementary setting to examine HVR2 evolution due to the physical isolation of freshwater bodies from one another in contrast to interconnected marine systems. These conditions provide a natural laboratory to study HVR2 dynamics under restricted gene flow, reduced recombination rates, and population bottlenecks. This study advances the characterisation of freshwater SAR11 HVR2 through the single-cell genomics analysis of Lake Biwa (Japan) and Lake Zurich (Switzerland) across multiple sampling years (2022 and 2024). Among available SAG technologies, the SAG-gel platform, which encapsulates each cell in an agarose gel bead, has been validated for seawater (Nishikawa et al., 2022) and freshwater samples (Okazaki et al., 2025) and was utilised in this project. The generated data constitute the largest single-cell genomic data for freshwater SAR11 HVR2 to date. In this study, we characterise the genomic architecture, gene content, and diversity of 176 *Fontibacterium* HVR2s extracted from Lake Biwa (2022, 2024) and Lake Zurich (2024), assess if HVR2 content stratifies by lake or sampling year, and compare their diversity and functional content against characterised marine counterparts.

## MATERIALS AND METHODS

### Designation of bacteria

Based on updated nomenclature, “*Candidatus* Fonsibacter” is now designated as *Fontibacterium* (Fernandes et al., 2025). Throughout this manuscript, we employ this genus name *Fontibacterium*. For example, “*Candidatus* Fonsibacter ubiquis” is referred to as *Fontibacterium ubiquis*.

### Samples and datasets

We sorted and sequenced 1920 droplets collected in 2022 from Lake Biwa in a previous study (Okazaki et al., 2025) and additionally sequenced 1152 and 3072 droplets in 2024 from Lake Biwa and Zurich, respectively. Briefly, the Biwa 2022 samples were collected in the summer (29 June 2022) and winter (1 February 2023) from the epilimnion and hypolimnion waters (Okazaki et al., 2025). Biwa 2024 samples were collected in August 2024 at 5m depth at a pelagic station (35°13909.50N, 135°59944.70E), and Zurich 2024 samples were collected in May 2024 from the epilimnion and hypolimnion near the Limnological Station Kilchberg (47°19′15.2″ N, 8°33′32.0″ E). All sampled droplets were sorted and sequenced via the SAG-gel technology (Nishikawa et al., 2022). Sequencing, read quality control and assembly of each SAG were performed in line with the methods of Biwa 2022 (Okazaki et al., 2025). The Biwa 2022 dataset comprised four samples: LB2206 (summer) and LB2301 (winter), each sampled from the epilimnion (E) and hypolimnion (H) (Okazaki et al., 2025). The Biwa 2024 dataset was designated BI2408E_sc. The Zurich 2024 dataset contained samples from the ZH2405E (epilimnion) and ZH2405H (hypolimnion), followed by a numerical ID.

Marine SAR11 genomes were obtained from Haro-Moreno et al. (2025), comprising 806 SAGs and isolates. They were first filtered by the presence of complete HVR2 which had already been identified by the previous authors. Following this, Prokka v1.14.6 (Seemann, 2014) (--rawproduct --mincontiglen 200) was utilised to identify the location of 23S and 5S genes in order to extract the HVR2. Ultimately, 145 complete HVR2 sequences were retrieved from the marine dataset and combined with our freshwater sequences, yielding in total 321 HVR2 sequences.

### Quality control

The freshwater SAGs were preliminarily analysed with CheckM v1.2.2 (Parks et al., 2015) (lineage_wf, default settings) for contamination and completeness assessments, QUAST v5.2.0 (Gurevich et al., 2013) (default settings) for contig statistics, and GTDB-Tk v2.3.2 (Chaumeil et al., 2022) (default settings; database release214) to elucidate their taxonomy.

### Extraction and characterisation of freshwater HVR2

All SAGs were first screened for the presence of the 23S and 5S rRNA genes, where Prokka v1.14.6 (Seemann, 2014) (--rawproduct --mincontiglen 200) was utilised for gene annotation. BLASTN searches via BLAST+ v2.9.0 (Camacho et al., 2009) were performed against 23S and 5S sequences obtained from a reference *Fontibacterium* isolate LSUCC0530 (Henson et al., 2018), and complete contigs that contained both rRNA sequences were selected. The 5S rRNA search retained hits with ≥99% nucleotide identity and 100% query coverage per HSP, keeping the single best hit per contig (- perc_identity 99, -qcov_hsp_perc 100, -max_target_seqs 1), whereas the 23S rRNA search used the default BLAST+ parameters; following this, manual curation was carried out to restrict the 23S to 99% identity and 2500bp in order to prevent false positive matches. HVR2 sequences were then extracted from the region between the 23S and 5S rRNA gene locations on each contig based on the coordinates obtained from the BLAST search, with strand directionality recorded. As an additional quality control step, we removed sequences longer than 1Mbp, as these likely originated from the opposite end of the circular chromosome. HVR2s were retained from all SAGs with these criteria, regardless of their initial GTDB-Tk assignment. Consequently, 176 freshwater HVR2 were extracted (32 from Biwa 2022, 135 from Biwa 2024 and 9 from Zurich 2024). Information about the HVR2s and their corresponding SAGs of origin is summarised in Supplementary Table 1.

### Comparison of HVR2 sequences

Diversity among HVR2 was calculated in two complementary approaches: highly sensitive, amino-acid sequence-based S_G_ scores and high-resolution, nucleotide sequence-based average nucleotide identity (ANI). Pairwise normalised tBLASTx scores (S_G_ scores) between all freshwater HVR2 sequences were computed with ViPTreeGen v1.1.2 (Nishimura et al., 2017). From the (0 ≤ S_G_ ≤ 1) range, a score of 1 indicates identical sequences, whereas a score approaching 0 indicates no detectable alignment between the pair (Bhunchoth et al., 2016). Sequence alignment between pairs of HVR2 was visualised by the DiGAlign webserver (Nishimura et al., 2024). Following HVR2 extraction, the remainder of each SAG genome (i.e., the genome with the HVR2 region excluded) was retained for a separate whole-genome comparison. In order to ensure accuracy, only SAGs classified as *Fonsibacter* (now *Fontibacterium*) by GTDB-Tk were retained, with additional filtering steps for medium- and high-quality genomes based on the minimum information about a single amplified genome (MISAG) standards by Bowers et al. (2017). pyANI-plus (Pritchard et al., 2016) was used to compute an all-vs-all ANI matrix of the HVR2s and genomic regions with the ANIb method (pyani-plus anib) under default parameters. The combined marine and freshwater HVR2 dataset was also analysed and visualised as described above.

### Functional annotation

Protein sequences on HVR2 were predicted using Prodigal v2.6.3 with default parameters and annotated using eggNOG-mapper v2 (Cantalapiedra et al., 2021) against the eggNOG database, which provided COG categories, PFAM domains and KEGG Orthology (KO) assignments. Additionally, the proteins were annotated with DRAM v1.4.6 (Shaffer et al., 2020) which provided an additional carbohydrate- active enzyme (CAZy) family assignment through dbCAN (Yin et al., 2012). The protein sequences were clustered into orthogroups using OrthoFinder v2 with default parameters. The relative position of each gene in the OG was calculated as the ratio of the gene start position to total HVR2 fragment length using the data in Prodigal GFF output files. HVR2 sequences were oriented from the 23S to the 5S gene before gene prediction positions were compared, so relative position is comparable across datasets. A comprehensive gene characterisation table was compiled for all 13817 OrthoFinder-assigned proteins, integrating annotations from eggNOG-mapper, DRAM, KEGG pathway assignments for a targeted subset of orthogroups via BlastKOALA, and genomic coordinates from GFF output files (Supplementary Table 2).

### Phylogenetic analysis

For the 3 core OGs (OG0000000, OG0000001, OG0000011), protein sequences from the HVR2 dataset were aligned using MAFFT v7.526 (Katoh et al., 2002). Maximum likelihood phylogenetic trees were inferred with IQ-TREE v3.0.1 (Wong et al., 2026) with model selection and 1000 ultrafast bootstrap replicates. The best-fit substitution models were VT+F+G4 for OG0000000 and OG0000001, and MTINV+F+I+G4 for OG0000011. iTOL v6 (Letunic & Bork, 2024) was used to visualise and annotate the trees, with annotation rings for dataset, habitat, KEGG K-Number and copy number per HVR2.

### Identification of tRNA genes in freshwater HVR2

The tRNA genes within individual HVR2 sequences were identified with tRNAscan-SE v2.0.12 (Chan & Lowe, 2019) in bacterial scanning mode. The location of tRNA midpoints was calculated from the annotated start and end coordinates and used to compute their distance to the 23S and 5S rRNA gene boundaries. The strand orientation of each tRNA was obtained from the tRNAscan-SE output and compared to the rRNA orientation to classify if they were co-directional or antisense relative to the rRNA.

## RESULTS

### Identification of HVR2 in SAR11 SAGs

Of the SAGs obtained from the three freshwater datasets, 392 were found to originate from *Fontibacterium* (6.7%). Of these, 200, 113 and 79 were obtained from Lake Biwa 2024, Lake Biwa 2022, and Lake Zurich 2024, respectively. In total, 176 HVR2-containing contigs were obtained from the dataset based on the flanking 23S and 5S rRNA genes. The Biwa 2024 dataset contributed the largest number of HVR2s (n = 135), followed by Biwa 2022 (n = 32) and Zurich 2024 (n = 9) (Table 1). Taxonomic annotation by GTDB-Tk revealed that HVR2-containing SAGs from Lake Biwa across the years were classified as *Fontibacterium ubiquis* and species-unresolved *Fontibacterium*. The Biwa 2024 dataset contained a small number of HVR2 originating from SAGs annotated as the phyla Actinomycetota and Pseudomonadota. As these sequences contained the *Fontibacterium* rDNA boundaries and satisfied our HVR2 detection criteria (see Methods for details), we retained them as bona-fide *Fontibacterium* HVR2 sequences, which likely originated from droplets encapsulating multiple cells. HVR2-containing SAGs from Lake Zurich (2024) were classified as *Fontibacterium* sp000510845 and species-unresolved *Fontibacterium*, consistent with the distinct *Fontibacterium* lineage composition of this geographically different freshwater system (Fernandes et al., 2025).

**Table 1.** Extracted HVR2 sequence statistics by sampling dataset and taxonomic origin.

| **Dataset** | **Dominant *Fontibacterium* Species** | **Sequenced Droplet Count** | **HVR2 Count** | **HVR2 Average Length (kb)** | **Coding Density** | **tRNA number** | **Source** |
|---|---|---|---|---|---|---|---|
| Biwa (2022) | *Fontibacterium ubiquis*<br>*Fontibacterium* sp. | 1920 | 32 | 52 | 0.88-0.98 | 32 | Okazaki et al. (2025) |
| Biwa (2024) | *Fontibacterium ubiquis*<br>*Fontibacterium* sp. | 1152 | 135 | 42 | 0.88-0.98 | 138 | This study |
| Zurich (2024) | *Fontibacterium* sp000510845<br>*Fontibacterium* sp. | 3072 | 9 | 36 | 0.90-0.98 | 9 | This study |

### Diversity of Freshwater HVR2s

The length of HVR2 sequences exhibited considerable variability across both freshwater and marine datasets (Figure 1a). Freshwater HVR2s (n = 176) ranged from 9.6 kb to 77.2 kb (median = 43.8 kb), whereas marine HVR2s (n = 145) ranged from 10.4 kb to 88.2 kb (median = 46.3 kb). The freshwater HVR2 datasets exhibited broadly comparable coding densities (0.88 to 0.98). Pairwise normalised tBLASTx (S_G_) scores were computed for all 176 freshwater HVR2 sequences, yielding 15,400 pairwise comparisons (Figure 1b). The distribution of S_G_ was strongly right-skewed, showing a high concentration in the low-similarity range (<0.2). A majority of S_G_ scores (14,970/15,400; 97.2%) fell below the 0.2 threshold, indicating that the freshwater pairs generally share few homologous genes. In the intra-sample analysis targeting only Biwa 2024 HVR2 sequences, the majority (8741/9045; 96.6%) of pairwise comparisons indicated S_G_ < 0.2 (Fig S1), suggesting the co-occurrence of highly divergent sequences even within sympatric populations. We also observed the occurrence of HVR2 pairs showing high sequence similarity only in part of their entire lengths (Figure 1c). We then performed inter-dataset analysis (Figure S2) and revealed that freshwater HVR2 sequences do not cluster by dataset origin, consistent with the high intra-dataset variability described above. As a function of genomic ANI (Figure 2a), HVR2 gene sharing exhibited a graded relationship. HVR2 S_G_ scores rose sharply above the 99% genomic ANI threshold, suggesting that genomes of the same population from the same lake can share near-identical HVR2 gene content, consistent with the vertical inheritance of HVR2. Below this threshold, HVR2 S_G_ scores broadly remain low across a broad range of genomic ANI values, indicating that HVR2 gene content quickly diverges independent of the rest of the genome; this evolutionary independence is already apparent among closely related genomes at ∼97.5% ANI, corresponding to a peak for the intra-population similarity.

**Figure 1.**
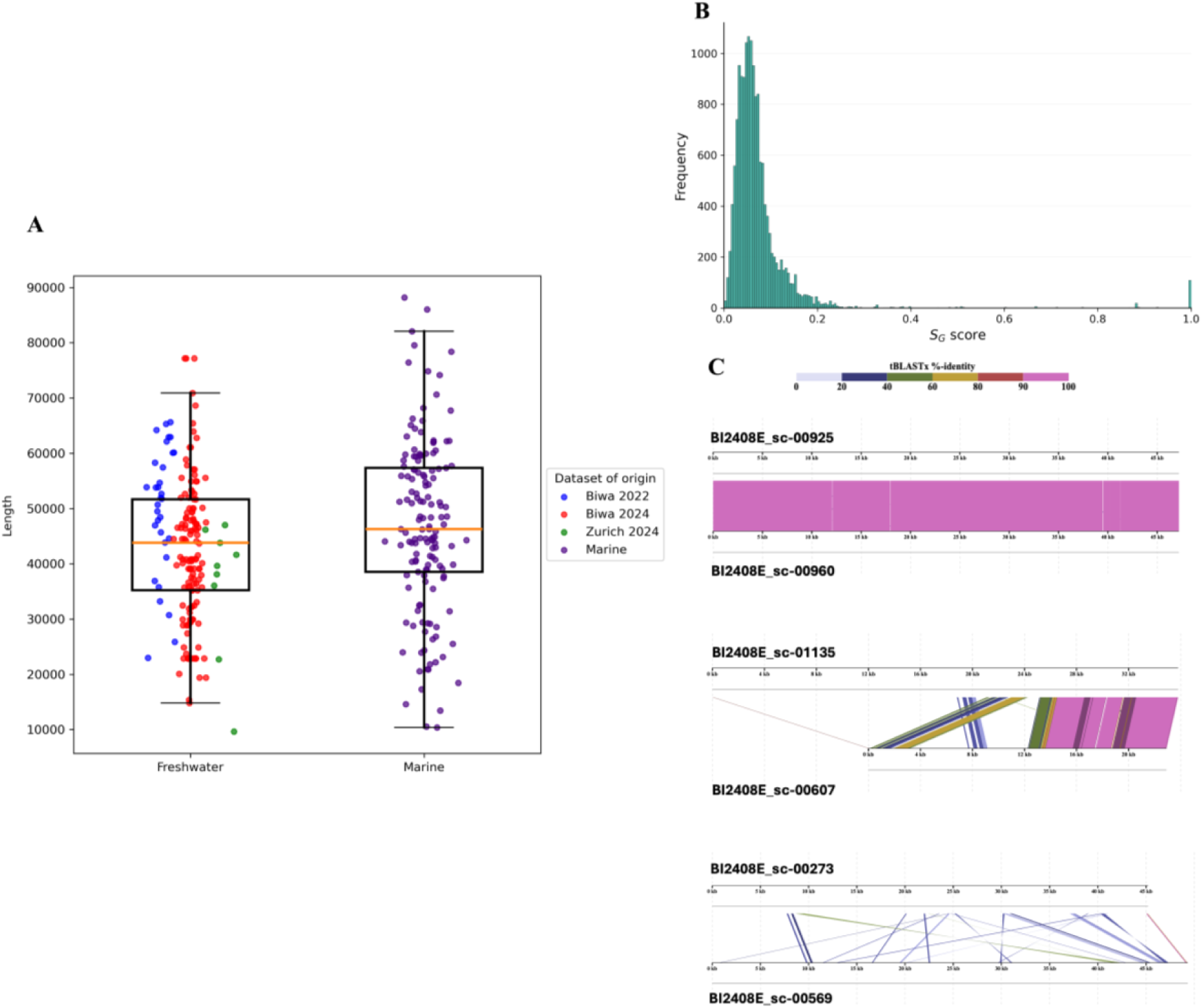
Length and normalised tBLASTx identity distributions in *Fontibacterium* HVR2. **(a)** Frequency distribution of HVR2 sequence lengths in freshwater and marine SAR11 datasets. The y- axis represents HVR2 length in nucleotide bases, and the x-axis represents the dataset origin. **(b)** Frequency distribution of pairwise normalised tBLASTx similarity (S_G_) scores among freshwater HVR2 sequences. The x-axis represents the S_G_ score (range 0.0–1.0), where higher values indicate greater shared homologous genes between HVR2 pairs, and the y-axis represents the number of pairwise comparisons. A total of 15,400 pairwise comparisons were performed across all 176 freshwater HVR2 sequences. The distribution is strongly right-skewed, with 97.2% of scores (14,970/15,400) falling below 0.2. **(c)** Adjacent panels show tBLASTx alignments for selected HVR2 pairs exhibiting various identities in descending order: high (BI2408E_sc-00925 and BI2408E_sc-00960), partial (BI2408E_sc-01135 and BI2408E_sc-00607) and low (BI2408E_sc-00273 and BI2408E_sc-00569) similarities.

**Figure 2.**
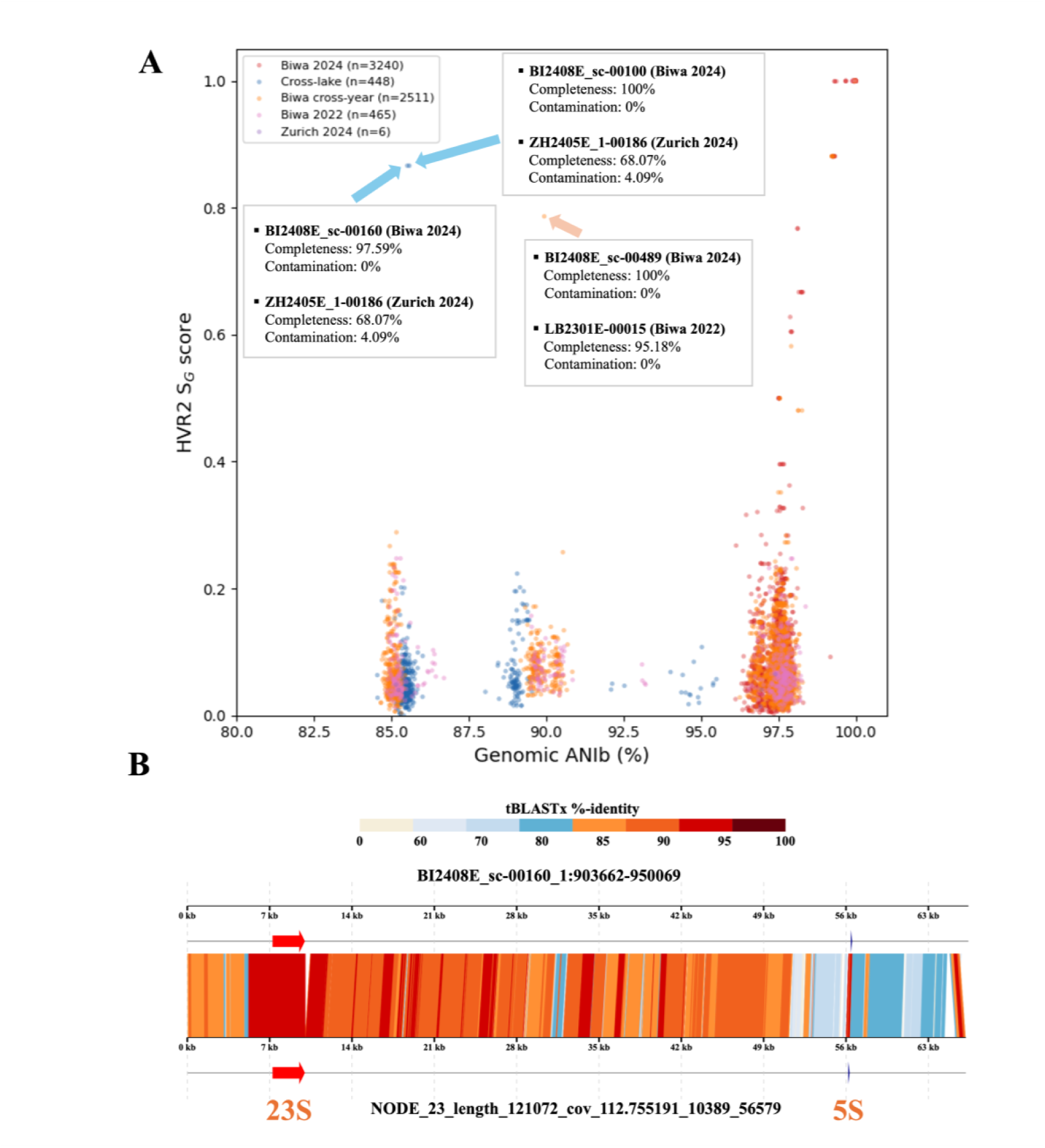
Comparison of Fontibacterium HVR2 S_G_ against genomic ANI and an intercontinental pair visualised through tBLASTx identity. **(a)** Scatterplot of HVR2 pairwise normalised tBLASTx (S_G_) scores against genomic (excluding HVR2) average nucleotide identity by BLAST (ANIb) for freshwater Pelagibacterales sequences. The x-axis represents genomic ANIb (%) and the y-axis represents HVR2 S_G_ score. Each point corresponds to a single pairwise comparison. S_G_ scores remain broadly low across a wide range of ANIb values, indicating that HVR2 gene content evolves independently of the surrounding genome, with a marked increase above approximately 99% genomic ANI. Blue arrows mark two cross-lake pairs with high S_G_ identity. Orange arrow marks a cross-year Lake Biwa pair with high S_G_ identity. **(b)** DiGAlign visualisation of a high-identity cross-lake HVR2 pair from Lake Biwa (contig BI2408E_sc-00160_1, SAG BI2408E_sc-00160) and Lake Zurich (contig NODE_23_length_121072_cov_112.755191, SAG ZH2405E_1-00186_S186). Pairwise tBLASTx alignment is shown, with colour intensity representing percentage amino acid identity. The 23S rRNA (red arrows) and 5S rRNA (blue arrows) flanking sequences demarcate the HVR2 boundaries. The S_G_ score for this pair was 0.8664, whilst the whole-genome ANIb excluding HVR2 was 85.5% (below the 95% species threshold). Median tBLASTx identity was 86.8% within HVR2 compared with 68.4% in the flanking regions.

There are two exceptions to this observation, in which HVR2s remain highly similar across species boundaries: one across continents and one within a single lake across sampling years. First, two HVR2 pairs between Lakes Biwa and Zurich showed gene synteny and remarkably high sequence similarity (Figure 2b). These were the only Biwa–Zurich pairs exceeding an S_G_ threshold of 0.5 (BI2408E_sc- 00160 and BI2408E_sc-00100 versus ZH2405E_1-00186_S186; S_G_ 0.866). The two Lake Biwa SAGs involved (BI2408E_sc-00160 and BI2408E_sc-00100) are themselves near-clonal (HVR2 S_G_ of 1.0; genomic ANI of 99.9%), so these two comparisons virtually represent a single cross-lake pair. The genomic ANIb between the Biwa–Zurich pair was 85.5%, below the 95% species threshold, consistent with the dominance of different *Fontibacterium* species in the two lakes as mentioned above. In contrast, the HVR2 S_G_ score was 0.866 and the HVR2 ANIb was 92.5%, indicating the strong conservation of gene content and synteny within this HVR2 pair despite a higher level of genome-wide divergence. Indeed, whole-contig alignment showed that the regions flanking HVR2 were considerably less similar than HVR2 itself (Figure 2b), with the median tBLASTx alignment identity in the flanking regions being 68.4%, compared with 86.8% within HVR2. Second, a similar case was observed between two Lake Biwa SAGs from different sampling years (BI2408E_sc-00489 and LB2301E-00015), which shared a genomic ANIb of 89.9%, below the species threshold, consistent with their GTDB-Tk classification: sc-00489 is *Fontibacterium ubiquis*, whereas LB2301E-00015 is a *Fontibacterium* lineage not assigned to a species. Despite this, their HVR2 S_G_ score was 0.787, whereas pairs at this level of genomic ANI typically have S_G_ scores near 0.06. The HVR2 ANIb identity (92.9%) was likewise higher than the genomic ANIb. In this pair, the HVR2 therefore remained more conserved than the rest of the genome.

### Comparison of HVR2 between Freshwater and Marine SAR11

Extended comparative analysis among marine and freshwater HVR2 sequences (Figure 3) revealed that most pairs showed low similarity (S_G_ < 0.2), consistent with the overall low gene-level conservation observed within the freshwater dataset alone. Within-habitat S_G_ score distributions revealed that a higher proportion of freshwater HVR2 pairs scored above 0.5 (165 pairs, 1.1%) than marine ones (6 pairs, 0.06%) (Figure 4). Consistently, none of the 25,520 cross-habitat (freshwater–marine) HVR2 pairwise comparisons exceeded the S_G_ threshold of 0.5 (mean S_G_ = 0.063, median = 0.055). Nevertheless, marine HVR2 sequences were clustered together with freshwater ones instead of forming habitat-specific clusters as seen in SAR11 species trees, and a small number of relatively high-scoring cross-habitat HVR2 pairs were detected (19 pairs at S_G_ > 0.3). The maximum cross-habitat S_G_ was 0.45, with all three top hits involving the same marine contig (contig ID CACKAY010000001.1_154806-193321, SAG ID GCA_902598265.1, recovered from the Bermuda Atlantic Time-series Study) paired against three different freshwater genomes – one from Lake Biwa 2022 and two from Lake Biwa 2024. This marine contig was taxonomically assigned to *Pelagibacter* through GTDB-Tk (*Pelagibacter* sp902616725), confirming that it belongs to a marine SAR11 lineage instead of freshwater *Fontibacterium*.

**Figure 3.**
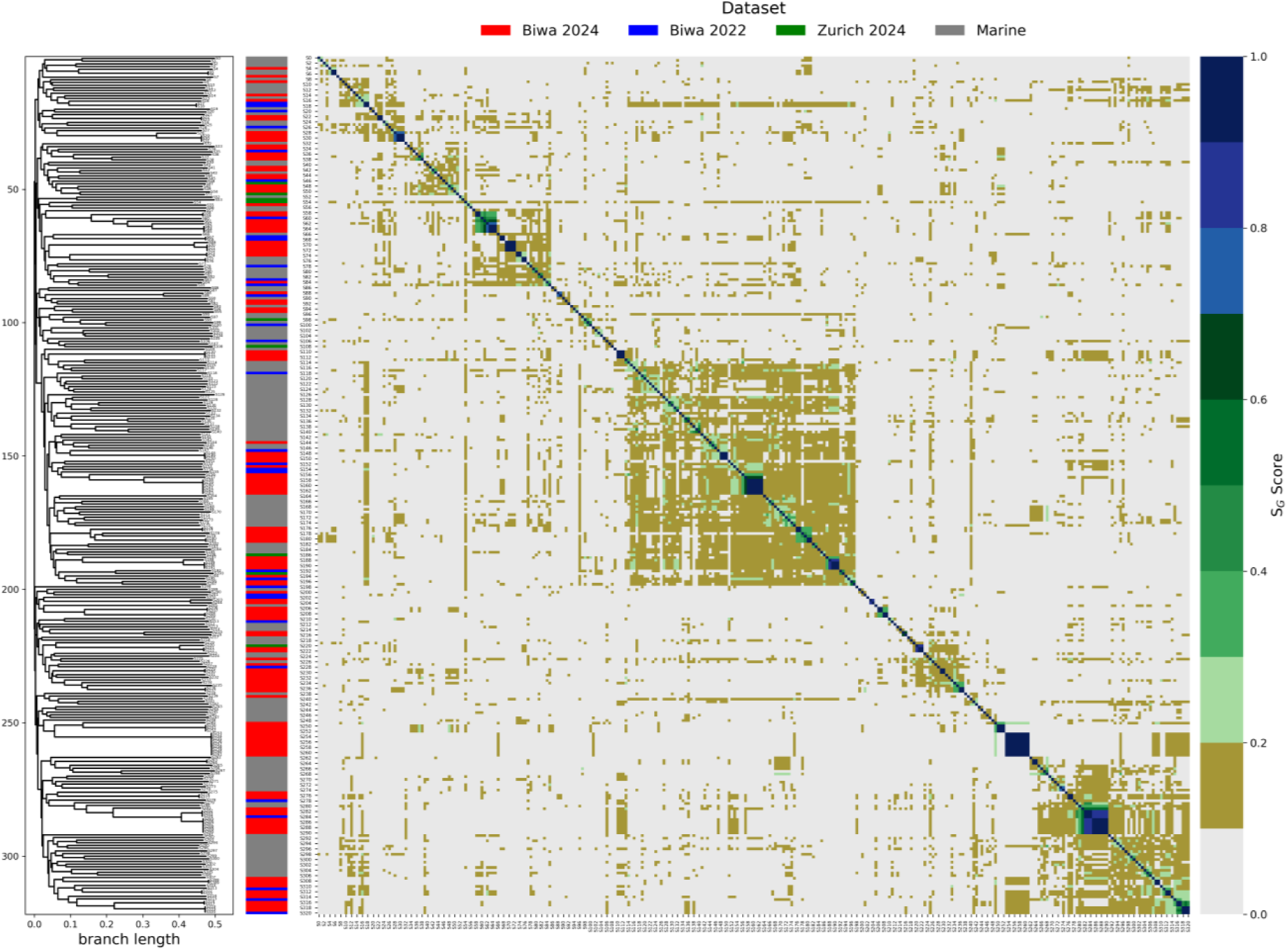
Pairwise S_G_ score heatmap of combined marine and freshwater HVR2 sequences. Normalised tBLASTx (S_G_) scores were computed for all pairwise comparisons between marine (n = 145) and freshwater (n = 176) HVR2 sequences. Colour intensity represents S_G_ score, with lighter shading indicating lower shared gene content. The dominant pattern is a broad low-similarity block (S_G_ < 0.2), with rare high-similarity pairs (S_G_ > 0.8). Marine and freshwater HVR2 sequences are interspersed rather than forming habitat-specific clusters.

**Figure 4.**
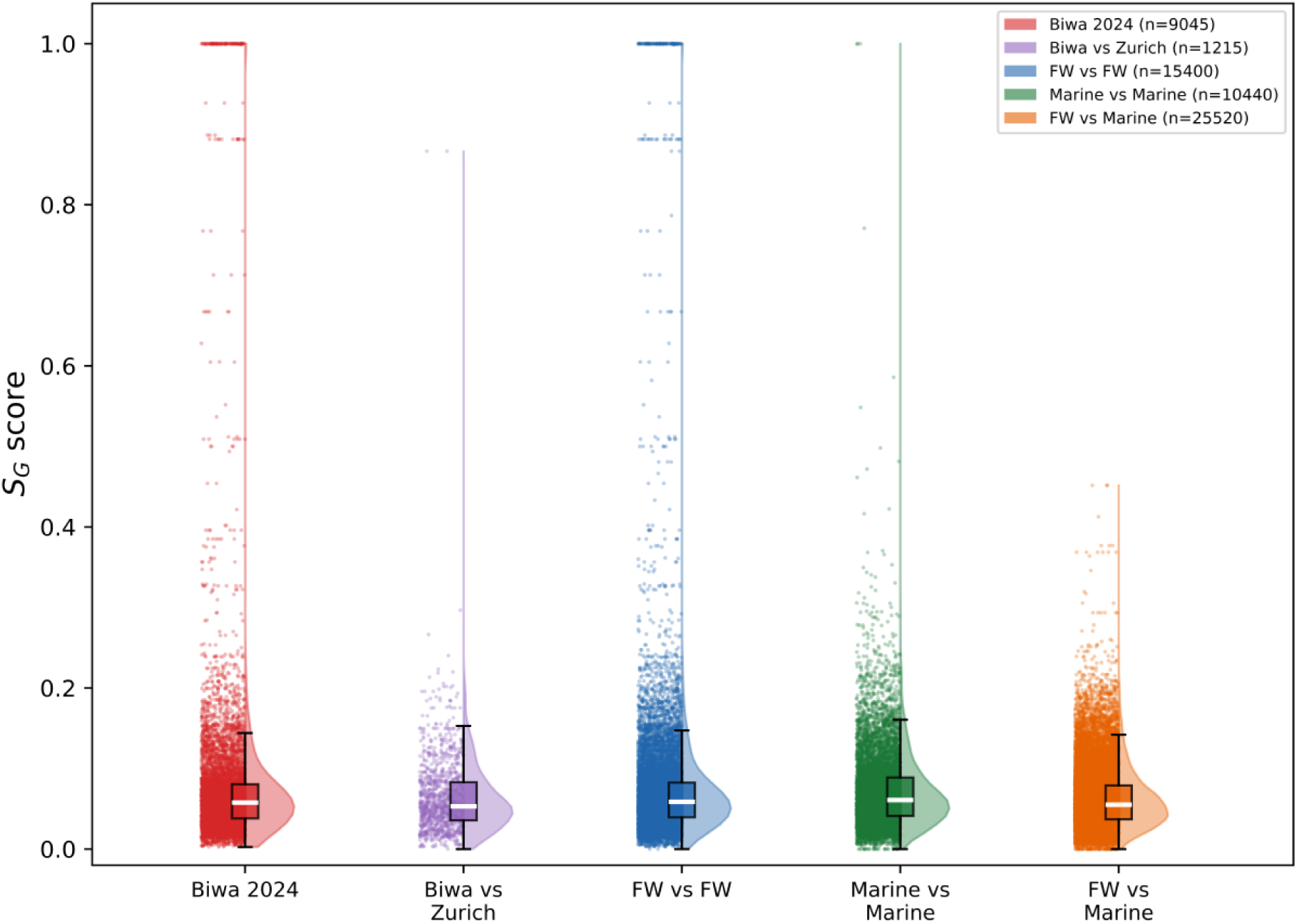
Comparison of pairwise S_G_ score distributions across sampling datasets and habitat types. Distributions are shown for within-freshwater, within-marine, cross-lake (Biwa–Zurich), and cross-habitat (freshwater–marine) pairwise HVR2 comparisons. Cross-habitat comparisons (n = 25,520 pairs) yielded a mean S_G_ score of 0.063 (median = 0.055), with no pairs exceeding 0.5. Within- freshwater comparisons produced 165 pairs above S_G_ = 0.5, whereas only 6 within-marine pairs exceeded this threshold. Only two cross-lake (Biwa–Zurich) pairs exceeded SG = 0.5.

### Functions Encoded in HVR2

Functional annotation of marine and freshwater HVR2 genes demonstrated that the dominant functional category across datasets was cell wall and membrane biogenesis (COG category M) among the 84.9% of genes that could be annotated (Figure 5a). Other functional categories included carbohydrate transport and metabolism, which may contribute to cell-surface polysaccharide modification. Of the 271 gene families clustered by OrthoFinder, 209 (77.1%) were shared between marine and freshwater datasets, indicating extensive cross-habitat functional overlap (Figure S3). A subset of 73 gene families was present in all four datasets, whereas 34 and 28 were exclusive to the freshwater and marine datasets, respectively. More than half the gene families (8/14, 57.1%) present in more than 50% of all HVR2 sequences (Table 2) were annotated as transferases acting on diverse substrates, including nucleotide sugars. Notably, three gene families were present in more than 90% of all HVR2 sequences (Figure 5b) and were designated as core gene families: OG0000000 (98.8% conserved; GDP-fucose biosynthesis / GDP-mannose 4,6-dehydratase), OG0000001 (98.4%; glycosyltransferase family 2), and OG0000011 (91.9%; ABC transporter transmembrane region). The relative positions of the three core gene families within HVR2 differed by family (Figure S4). OG0000001 was skewed towards the 5S rRNA end (mean relative position 0.69), with 694 (63.7%) genes located in the final third of HVR2. In contrast, OG0000000 and OG0000011 showed no clear preference for either rRNA end and were distributed throughout HVR2. Phylogenetic analysis of the three core genes revealed that marine and freshwater sequences do not form separate clades but are intermixed in each tree (Figure S5a, b, c). OG0000011 (Figure 5c) was notable for its low copy number per HVR2, with only 1 to 2 copies present in each HVR2 encoding this gene family (single copy in all but 12/295 HVR2 sequences), whereas OG0000000 and OG0000001 were more numerous, typically present in the range of 3 to 12 copies in each HVR2.

**Figure 5.**
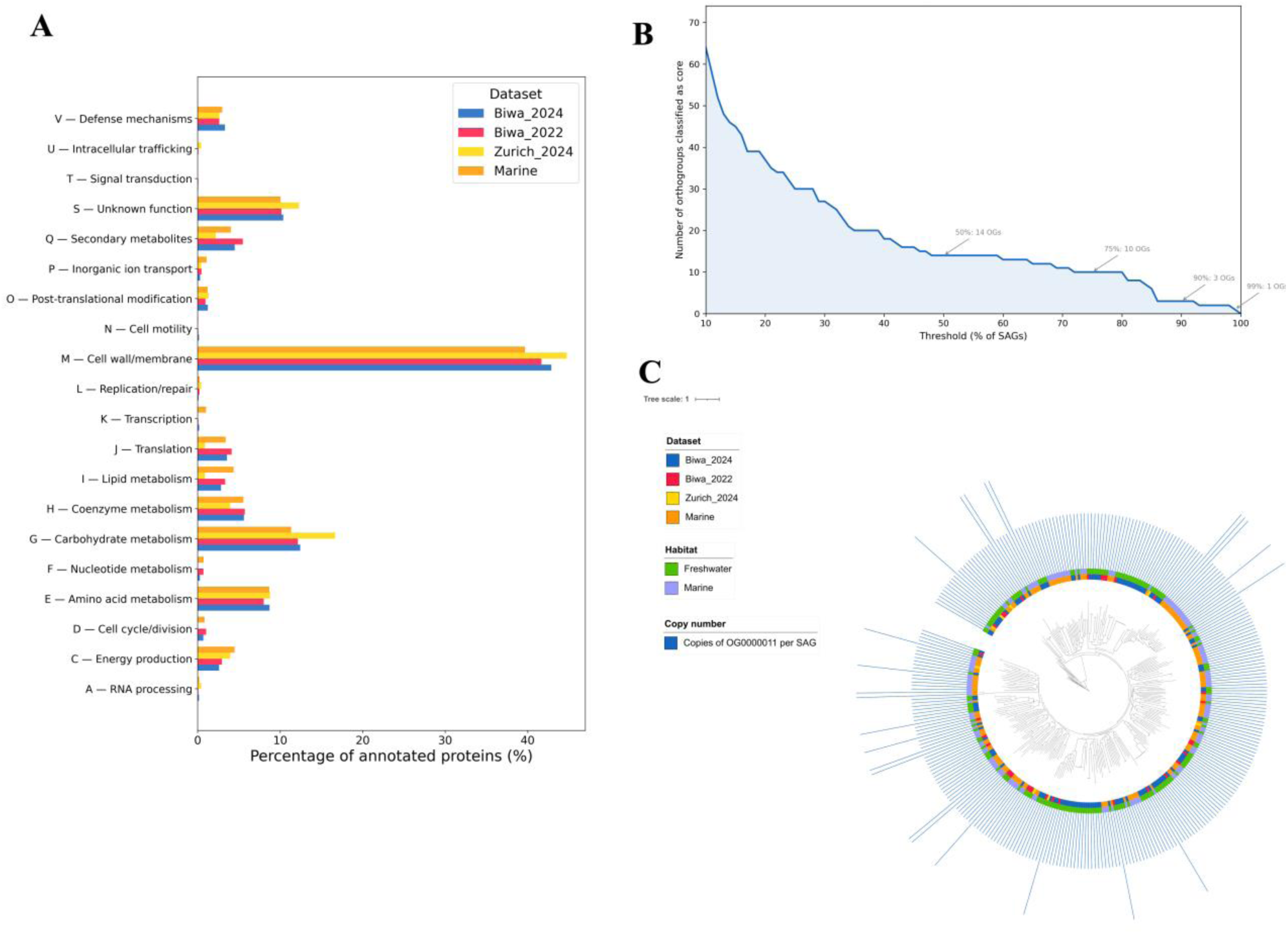
Functional annotation of marine and freshwater HVR2 proteins. **(a)** Distribution of Clusters of Orthologous Groups (COG) functional categories among predicted HVR2 proteins in marine and freshwater datasets. Functional annotation was performed using eggNOG-mapper. The x-axis shows COG functional categories, and the y-axis shows the number of annotated proteins per category. Category M (cell wall, membrane, and envelope biogenesis) was the dominant functional category in both habitats, consistent with the established role of HVR2 in surface polysaccharide biosynthesis. 84.9% of the proteins were annotated and displayed in the figure, whereas the unannotated 15.1% are not. **(b)** Conservedness of gene families across marine and freshwater HVR2 sequences. The x-axis represents the threshold of conservation of individual gene families (% of HVR2 sequences containing the gene family), and the y-axis represents the number of gene families. Gene families present in more than 50% of all HVR2 sequences (listed in Table 2) were predominantly annotated as transferases. Three core gene families exceeded 90% conservedness: OG0000000 (98.8%), OG0000001 (98.4%), and OG0000011 (91.9%). **(c)** Maximum likelihood phylogenetic tree of OG0000011 across marine and freshwater HVR2 sequences. The tree was constructed using IQ-TREE 3, with sequences aligned by MAFFT. Marine and freshwater sequences are intermixed throughout each tree rather than forming habitat-specific clades. OG0000011 (ABC transporter transmembrane region) was notable for its low copy number (1–2 copies per HVR2).

**Table 2.** List of annotated orthogroups up to 50%.

| OG_ID | Gene_description (most abundant annotation) | Distinct KO | Conservedness (%) |
|---|---|---|---|
| OG0000000 | Catalyzes the two-step NADP-dependent conversion of GDP- 4-dehydro-6-deoxy-D-mannose to GDP-fucose, involving an epimerase and a reductase reaction / Catalyzes the conversion of GDP-D-mannose to GDP-4- dehydro-6-deoxy-D-mannose [MIXED] | True | 98.8 |
| OG0000001 | Glycosyltransferase like family 2 / Glycosyl transferase family 2 [MIXED] | True | 98.4 |
| OG0000011 | ABC transporter transmembrane region / ATPases associated with a variety of cellular activities [MIXED] | True | 91.9 |
| OG0000007 | NeuB family / acid synthase [MIXED] | True | 85.7 |
| OG0000013 | phospho-N-acetylmuramoyl-pentapeptide-transferase activity / Glycosyl transferase family 4 [MIXED] | True | 85 |
| OG0000002 | Methyltransferase domain / methyltransferase [MIXED] | True | 85.0 |
| OG0000012 | O-antigen | False | 83.8 |
| OG0000003 | Glycosyl transferases group 1 / transferase activity, transferring glycosyl groups [MIXED] | True | 83.5 |
| OG0000005 | Belongs to the DegT DnrJ EryC1 family | False | 80.1 |
| OG0000004 | Cytidylyltransferase / Aminotransferase class-III [MIXED] | True | 79.8 |
| OG0000006 | Polysaccharide biosynthesis protein / Catalyzes the conversion of GDP-D-mannose to GDP-4- dehydro-6-deoxy-D-mannose [MIXED] | True | 71.7 |
| OG0000008 | Nucleotidyl transferase / D,D-heptose 1,7-bisphosphate phosphatase [MIXED] | True | 67.9 |
| OG0000010 | Methyltransferase FkbM domain / Specifically methylates the adenine in position 37 of tRNA(1)(Val) (anticodon cmo5UAC) [MIXED] | True | 64.5 |
| OG0000009 | Acetyltransferase (GNAT) domain / Attaches a formyl group to the free amino group of methionyl-tRNA(fMet). The formyl group appears to play a dual role in the initiator identity of N-formylmethionyl-tRNA by promoting its recognition by IF2 and preventing the misappropriation of this tRNA by the elongation apparatus [MIXED] | True | 59.5 |

### tRNA Genes as Putative Anchors for Recombination

At least one tRNA gene was identified in every freshwater HVR2 sequence examined (n = 176). All tRNAs were classified as initiator methionine (fMet), yielding 179 tRNA genes in total across the three datasets. Most HVR2s (173/176, 98.3%) contained exactly one tRNA gene, with exceptions from 3 HVR2 sequences in Biwa 2024, where a second tRNA-Met gene was detected. The genes were concentrated near the 5S rRNA gene, with 76.5% lying in the last fifth of the HVR2 (relative position 0.8-1.0, Figure S6a) and 100 genes (56%) were within 300 bp of the 5S rRNA gene, with 90 of them concentrated between 145-160 bp from the 5S rRNA gene. Most genes (85%, 152/179) were on the opposite strand of the rRNA operon.

## DISCUSSION

We recovered 176 complete *Fontibacterium* HVR2 sequences from SAGs collected from two geographically separated lakes (Lake Biwa, Japan and Lake Zurich, Switzerland) across multiple sampling years. The scale of the dataset is comparable with those established for marine HVR2 (Haro- Moreno et al., 2025), allowing for cross-habitat comparative analyses of the variable regions. As a result, freshwater HVR2s exhibit extreme diversity within a sympatric population similar to their marine counterparts, with the region evolving largely independently of the rest of the genome. The comparable lengths between freshwater and marine HVR2 (respective medians of 43.8 kb and 46.3 kb) indicate that HVR2 size is generally conserved across SAR11 (Figure 1a), consistent with previously reported architectural conservation in marine subclades (Haro-Moreno et al., 2025). The strongly right-skewed S_G_ distribution, with 97.2% of pairwise comparisons scoring below 0.2 (Figure 1b), demonstrates that few homologous genes are shared between HVR2 pairs, even within the same sampling location and time point. This low level of conservation was also seen within a sympatric population from the same time in the same lake (Figure S1). This observation mirrors the extensive intra-population HVR2 diversity documented for marine SAR11 (Molina-Pardines et al., 2025). Nevertheless, a small minority of freshwater HVR2s can be highly conserved across species, years, and lakes, including two high- similarity cross-continental pairs (Figure 2). Moreover, we found conserved gene families, including three nearly universal ones, not only among freshwater, but across freshwater and marine HVR2s. The initiator methionine tRNA genes enriched near the 5S rRNA gene region suggest that the tRNA genes may play a role in homologous recombination.

The intriguing coexistence of the diversity and conservation in HVR2 can be explained through different scenarios depending on the genomic relatedness of the bacteria. Intraspecies genomic ANI was concentrated around the 97.5% threshold rather than 100%, indicating that the sampled population is not clonal (Figure 2a). At a closer level of relatedness, near-identical HVR2 (S_G_ ≈ 1.0) sequence is shared almost exclusively by pairs with genomic ANI above 99%, indicating that the HVR2 can be co- inherited with the rest of the genome through clonal vertical transmission, leaving the genetic divergence of the HVR2 similar to the level of its genomic background. In contrast, below this threshold, the HVR2 S_G_ score was already low, suggesting that HVR2 diversifies much faster than the rest of the genome within a population. Despite this hypervariability, there were a few highly similar (S_G_ > 0.7) HVR2 pairs across the dataset (Figure S2), suggesting that HVR2 evolves independently of its surrounding genomic context. A notable example was found from Lake Biwa, in which two highly similar HVR2s (S_G_ = 0.787) recovered across different sampling years shared a genomic ANIb of only 89.9%. Since these SAGs originate from the same lake, this points to within-lake conservation of an HVR2 genotype across the species boundary and time. Another notable instance was the occurrence of a high identity (S_G_ = 0.866) HVR2 pair with a conserved genetic synteny between Lake Biwa and Lake Zurich (Figure 2b). The lack of hydrological connection between the lakes may make horizontal gene transfer implausible. However, it is known that the same species of microbes can be detected in geographically distant lakes (Fernandes et al., 2025, Hoetzinger et al., 2024), which implies the rare possibility of intercontinental genetic exchanges. This, combined with the distinct species assignments of their SAGs (*Fontibacterium ubiquis* and *Fontibacterium sp000510845* for Lakes Biwa and Zurich respectively), implies that the similarity of the pair may reflect an indirect HVR2 transfer or parallel acquisition of related HVR2 from unknown sources. This interpretation is further supported by the lower similarity of their flanking regions (median flank tBLASTx identity: 68.4%). The existence of similar HVR2 between distinct species in geographically separate populations mirrors another case of HVR2 conservation described by Haro-Moreno et al. (2025), where distantly related species (∼85% AAI) could share similar HVR2s, albeit in the minority of cases. Finally, local blocks of conserved regions could be found in otherwise divergent HVR2s (Figure 1c). This result suggests the possibility of recurrent partial insertions or recombinations within HVR2 instead of the replacement of entire HVR2 regions through recombination, implying an additional mechanism of HVR2 diversification. Overall, our results suggest the existence of a pool of HVR2 sequences that evolve through mutual genetic exchanges, independently of the rest of the genome.

Functional annotation of the combined marine-freshwater HVR2 dataset revealed substantial conservation of their gene functions. Cell wall, membrane, and envelope biogenesis (COG category M) is the dominant functional category in both habitats, consistent with the hypothesised role of HVR2 in surface polysaccharide biosynthesis (Grote et al., 2012). The enrichment of functional categories mirrors the previous analysis of marine SAR11 HVR2, where strains carry largely different HVR2 genes that nonetheless encode similar biosynthetic functions, so the cell surface is rebuilt using interchangeable (e.g., paralogous or non-homologous) gene sets rather than by a shared set of orthologues (Molina-Pardines et al., 2025). Our findings show that this observation extends across the marine-freshwater divide: although freshwater and marine SAR11 HVR2 largely possess different individual HVR2 genes, they encode enzymes of the same functional categories. Moving from broad functional categories (COG) to individual gene families (OG), the families shared between habitats were predominantly transferases acting on diverse substrates (Figure 5a; Table 2), reinforcing that the conserved functional core of HVR2 is cell-surface carbohydrate modification. Beyond this shared core, a minority of families were restricted to a single habitat, with distinct sets exclusive to the marine and to the freshwater datasets (Figure S3). Their existence suggests that, alongside the conserved core gene families, each HVR2 also maintains a small habitat-specific complement of gene families.

Three gene families were nearly universally conserved across freshwater and marine HVR2s: OG0000000, OG0000001, and OG0000011 (Figure 5c, Figure S5). OG0000000 is mostly annotated as enzymes involved in the biosynthesis of three related activated sugar donors – GDP-fucose, GDP- mannose and dTDP-rhamnose, all of which are needed for the synthesis of O-antigen and cell-surface polysaccharides in Gram-negative bacteria (Lerouge & Vanderleyden, 2002; Reeves et al., 1996).

OG0000001 is primarily annotated as glycosyltransferase family 2, which catalyses the transfer of sugars onto growing polysaccharide chains (Lairson et al., 2008), consistent with the cell-surface modification characteristics. OG0000011 (Figure 5c) primarily encodes an ABC transporter transmembrane region and is notably present at low copy number (1–2 copies per HVR2), though the reason for this occurrence remains unclear. In Gram-negative bacteria, ABC transporters are commonly part of the core machinery for the transport of O-antigen and cell-surface polysaccharides (Cuthbertson et al., 2010). The placement of two of the three gene families at opposite ends of the HVR2 (OG0000001 at the 5S end, OG0000011 at the 23S end) suggests that gene content may be positionally structured within the region, rather than randomly distributed. Maximum-likelihood phylogenetic analysis of the three OGs demonstrated that marine and freshwater HVR2s are intermixed and do not separate in the trees, indicating that these core functions, but not necessarily with the same functions, are important across habitats, alluding to their significance for a wide range of SAR11 (Figure S5). Together, these findings establish that marine and freshwater HVR2 share functional gene families that predominantly comprise cell-surface polysaccharide biosynthesis machinery.

At least one initiator methionine (fMet) tRNA gene was present in every examined freshwater HVR2 (n=176), yielding 179 tRNA genes in total (Figure S6a). tRNA genes are well-documented integration anchors for tyrosine-integrase-mediated genomic islands across bacterial lineages (Williams, 2002), including in streamlined marine cyanobacteria such as *Prochlorococcus* (Liu & Zhu, 2010). Methionine tDNAs have been established as integration hotspots in other bacteria, including the *Klebsiella pneumoniae* met56 tRNA-Met locus (Van Aartsen et al., 2012) and the *Vibrio cholerae* VC0516.1 locus underlying Vibrio Seventh Pandemic Island II (Dziejman et al., 2001; Murphy & Boyd, 2008). The universal presence of at least one fMet tRNA gene across freshwater HVR2s is therefore compatible with a role in site-specific integration. It was previously suggested that the conserved flanking rRNA anchors may enable each HVR2 to be transferred as a cassette (Haro-Moreno et al., 2025). While the conserved rRNA boundaries bounding every HVR2 may be the primary sites of homologous recombination, the concentration of methionine tDNAs near these boundaries raises the possibility of a complementary mechanism in the integration of the region, albeit untested. This is especially pronounced from the bias towards antisense orientation and concentration at the 5S region, which departs from a simple expectation of random placement, though its cause remains unresolved. These findings are also consistent with Haro-Moreno et al. (2025), who identified the presence of valine and methionine tRNAs near the 5S end of the marine HVR2. Overall, these findings indicate a role for tRNAs as anchors for insertion and recombination.

## CONCLUSION

The main conclusion of this study is paradoxical: while SAR11 HVR2 is extremely variable, it occasionally exhibits signs of evolutionary conservation beyond the phylogenetic and habitat divides. By integrating freshwater and marine HVR2 datasets into a single comparative analysis, this study establishes the features of HVR2 evolution that are shared across the SAR11 order. Notably, despite its extreme sequence divergence independent of overall genomic similarity, HVR2 remains constrained at both the sequence and functional composition levels across the marine-freshwater divide. This pattern, in which divergent genes still contribute to related phenotypes, appears to be a general feature of SAR11 HVR2. Together, our results position HVR2 as a functionally conserved yet evolutionarily dynamic region and raise novel questions to be resolved in future studies: How is HVR2 diversity generated and maintained? How large is its genetic diversity within a population and within ecosystems? What is the selective pressure responsible for their sequence and functional preservation? How are HVR2 genes transferred across species and habitat divides? As SAG sampling expands to more lakes and broader spatiotemporal scales, capturing the full extent of HVR2 diversity, research on freshwater SAR11 HVR2 will provide further valuable insights into the eco-evolutionary dynamics of microbial genomes and test whether the patterns observed here hold more broadly.

## ACKNOWLEDGMENTS

This work was supported by the Japan Society for the Promotion of Science (JSPS KAKENHI 21H05057, 22K15182 and 25K18161), the Japan Science and Technology Agency (JST FOREST JPMJFR2273 and ACT-X JPMJAX20BE), and the Collaborative Research Program of the Institute for Chemical Research, Kyoto University (2024-28). Computational resources were provided by the Supercomputer System of the Institute for Chemical Research, Kyoto University. Sample collection in Lake Biwa was supported by the Center for Ecological Research, Kyoto University, a Joint Usage/Research Center and the research vessel ‘Hasu’. S.A. was supported by the grants RPG_072023_02 (Swiss State Secretariat for Education, Research and Innovation) and PZ00P3_193240 (Swiss National Science Foundation). Y.O. was supported by the Kyoto University – University of Zurich ECR Mobility program. An AI tool was used for language proofreading and script refinement purposes, and all generated outputs were carefully reviewed and approved by the authors prior to their use in this study. We thank Jakob Pernthaler, Lucas Serra Moncadas, Cyrill Hofer, Alisa Shakurova, Liwen Zhang, Yukiko Goda and Tetsuji Akatsuka for their assistance in the field sampling.

## DATA AVAILABILITY

The nucleotide sequences of the HVR2 and SAGs are available at GenomeNet: https://www.genome.jp/ftp/db/community/FWSAG_HVR2/

## SUPPLEMENTARY TABLE LEGENDS

**Supplementary Table 1. Information on the 176 freshwater HVR2 sequences and their SAG of origin.** Each row contains an HVR2 and its corresponding SAG ID (32 from Biwa 2022, 135 from Biwa 2024, and 9 from Zurich 2024). The HVR2 refers to the region between the 23S and 5S rRNA genes. *HVR2 Info:* **HVR2_ID** refers to the HVR2 sequence ID (contig ID followed by the start and end of the HVR2 on that contig); **HVR2_Length** refers to the length of the HVR2 (bp); **Dataset**, Biwa 2022, Biwa 2024 or Zurich 2024; **HVR2_FASTA_Orientation** refers to the direction in which the HVR2 sequence is stored in the FASTA file (23S→5S or 5S→23S). *HVR2-Containing SAG Info:* **SAG_ID**, single-amplified genome identifier; **SAG_Length**, total assembly length (bp); **Completeness** and **Contamination,** obtained from CheckM estimates (%); **GC**, assembly GC content (%); **N50**, assembly N50 (bp); **classification**, GTDB taxonomy from GTDB-Tk (an empty species field, s, means no species was assigned); **num_16S**, **num_23S**, **num_5S**, numbers of rRNA genes in the assembly; **num_contigs**, number of contigs.

**Supplementary Table 2. Proteins predicted from the HVR2 sequences, with orthogroup and annotation.** Proteins were predicted with Prodigal from the 321 HVR2 sequences (176 freshwater, 145 marine) and grouped with OrthoFinder. One row per protein, ordered by SAG and then gene start. **Protein_ID** is the HVR2 file key and protein ID separated by | (the protein number follows the order along the stored sequence); **HVR2_FASTA_Orientation** refers to the direction in which the HVR2 is stored (as in Supplementary Table 1); **SAG_ID** is the corresponding SAG of origin; **Dataset,** Biwa 2022, Biwa 2024 or Zurich 2024; **Habitat,** Freshwater or Marine; **Orthogroup**, Orthogroups classified by OrthoFinder; **Conservedness_pct** is the percentage of the HVR2s containing the orthogroup; **Copy_number_per_HVR2** is the number of proteins of this orthogroup in each HVR2; **COG_category**, **PFAM_eggnog**, **PFAM_dram**, **eggNOG_description**, **Preferred_name**, **KEGG_ko_eggnog** are obtained from eggNOG-mapper and DRAM annotations (- or empty = none); **KEGG_ko_blastkoala** is obtained from KEGG orthology from BlastKOALA; **KEGG_ko_dram**, **DRAM_rank** (confidence, A highest to E lowest), **KEGG_hit_dram**, **CAZy_ids**, **CAZy_best_hit** are from DRAM annotations; **KEGG_pathway_broad** and **KEGG_maps** are from pathway and KEGG maps linked to the BlastKOALA KO; **Gene_start** and **Gene_end** are coordinates on the HVR2 sequence as stored in the FASTA; **Gene_orientation** is the gene strand (+ or −) relative to the stored HVR2 sequence.

---

**Figure S1.**
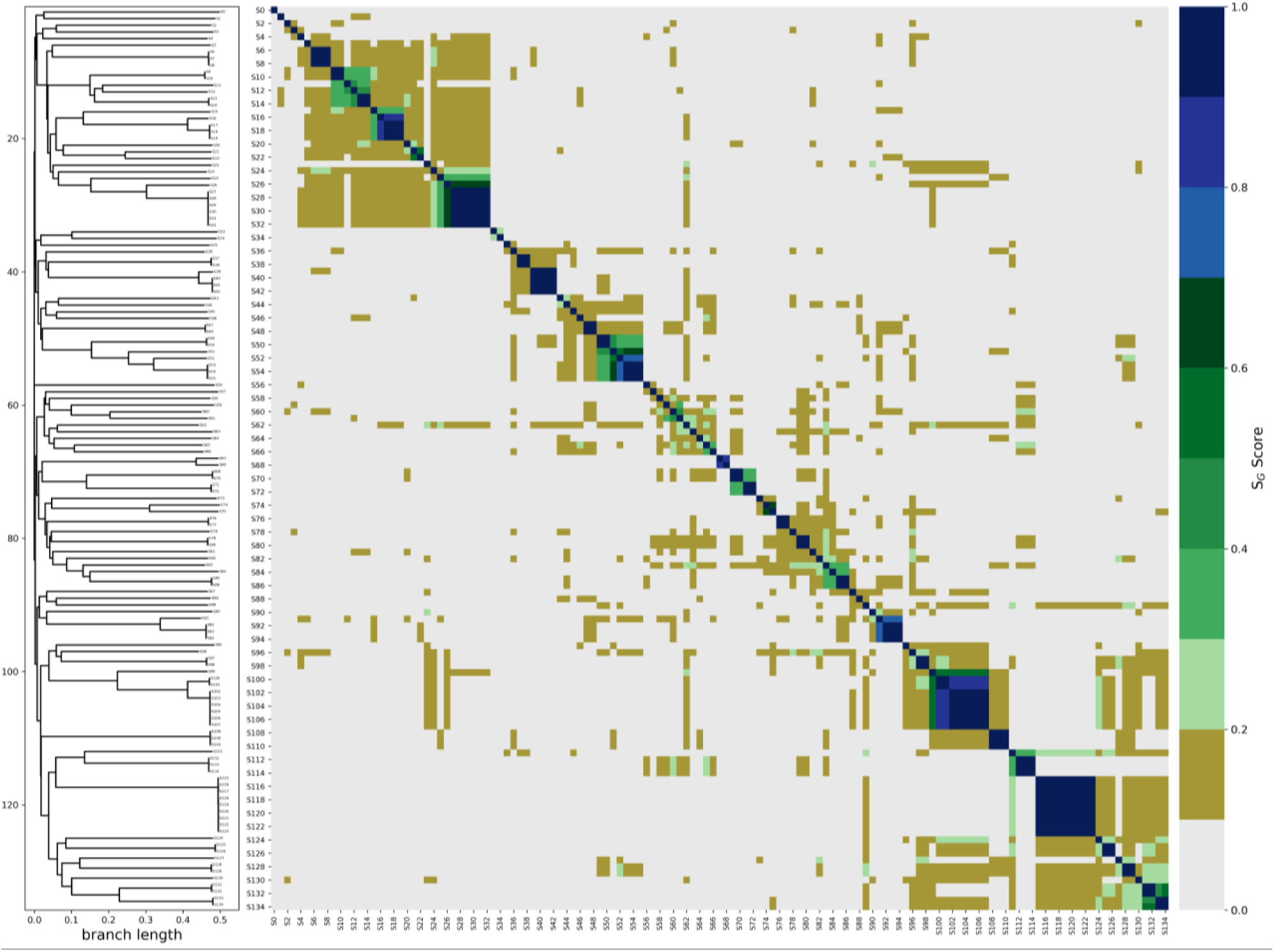
Pairwise similarity heatmap with UPGMA clustering of Lake Biwa 2024 HVR2 sequences with selected tBLASTx alignments. The heatmap displays pairwise S_G_ similarity scores among HVR2 sequences from the Biwa 2024 dataset, with colour intensity corresponding to the degree of shared gene content. The majority of pairwise comparisons (96.6%) fell below 0.2 S_G_, reflecting high intra-dataset HVR2 divergence.

**Figure S2.**
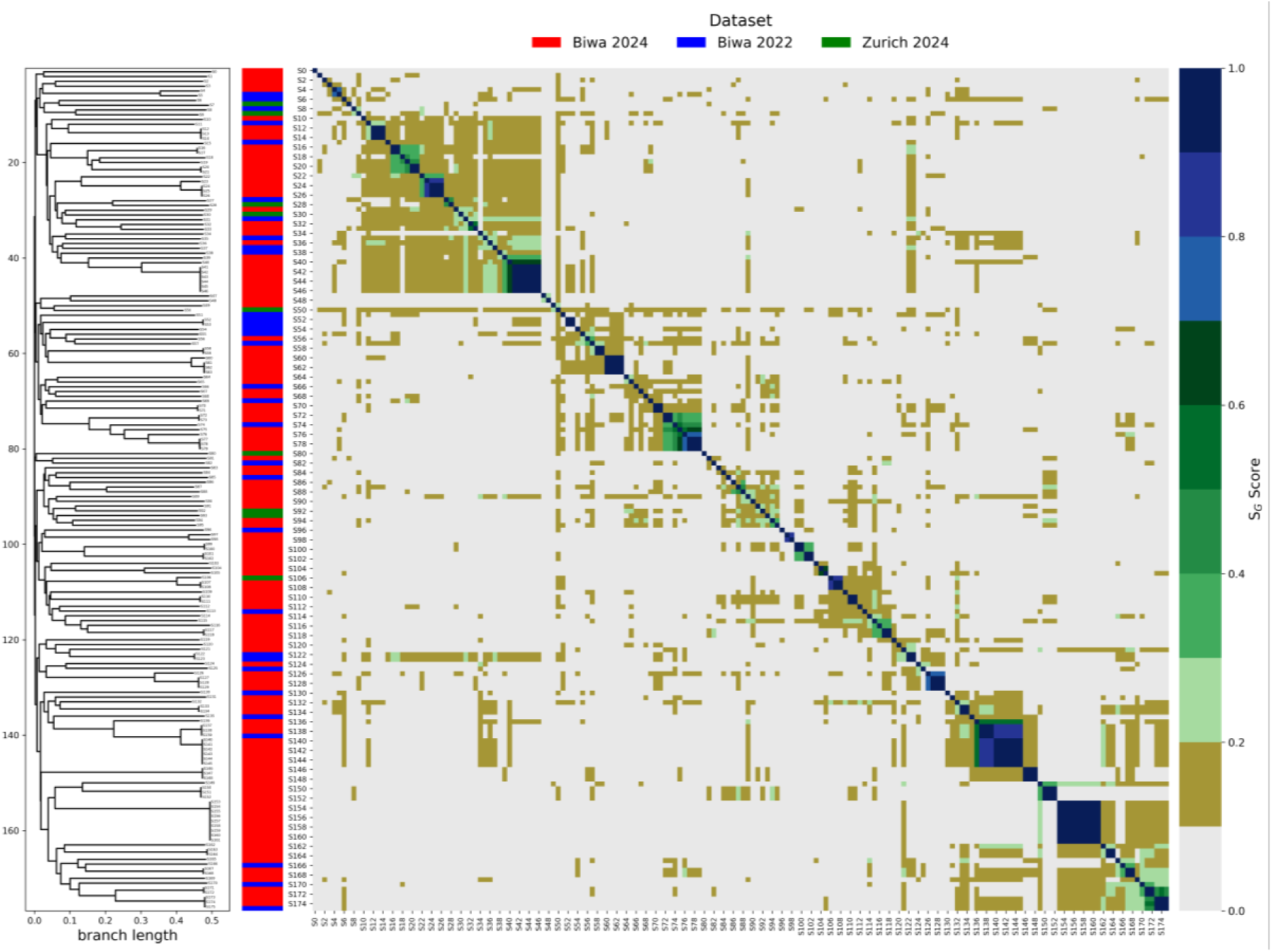
Pairwise normalised tBLASTx (S_G_) score heatmap with UPGMA clustering of combined freshwater HVR2 sequences across all datasets. HVR2 sequences from Lake Biwa and Lake Zurich datasets were combined, and pairwise S_G_ scores were computed. Colour intensity represents S_G_ score, with darker shading indicating higher shared gene content. A minority of pairs with S_G_ scores above 0.7 form clusters along the diagonal, indicating a small number of highly conserved HVR2 sequences shared across datasets.

**Figure S3.**
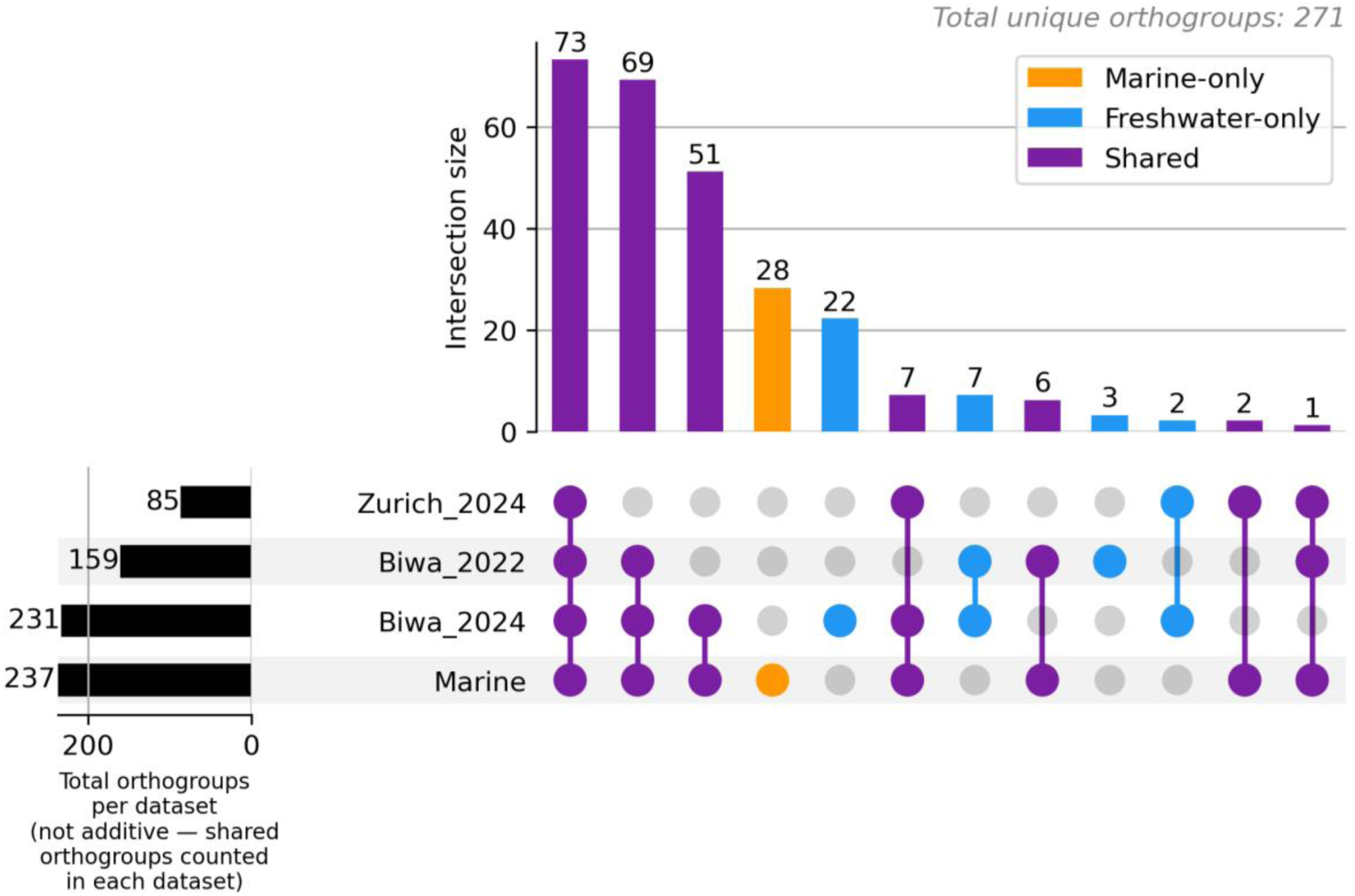
Distribution of gene families identified by OrthoFinder across marine and freshwater HVR2 datasets. A total of 271 gene families were identified from predicted HVR2 proteins, of which 209 (77.1%) were shared between marine and freshwater datasets. A core subset of 73 gene families was present across all four sampling datasets (Biwa 2022, Biwa 2024, Zurich 2024, marine). Of the 62 habitat-restricted gene families, 28 were exclusive to the marine dataset.

**Figure S4.**
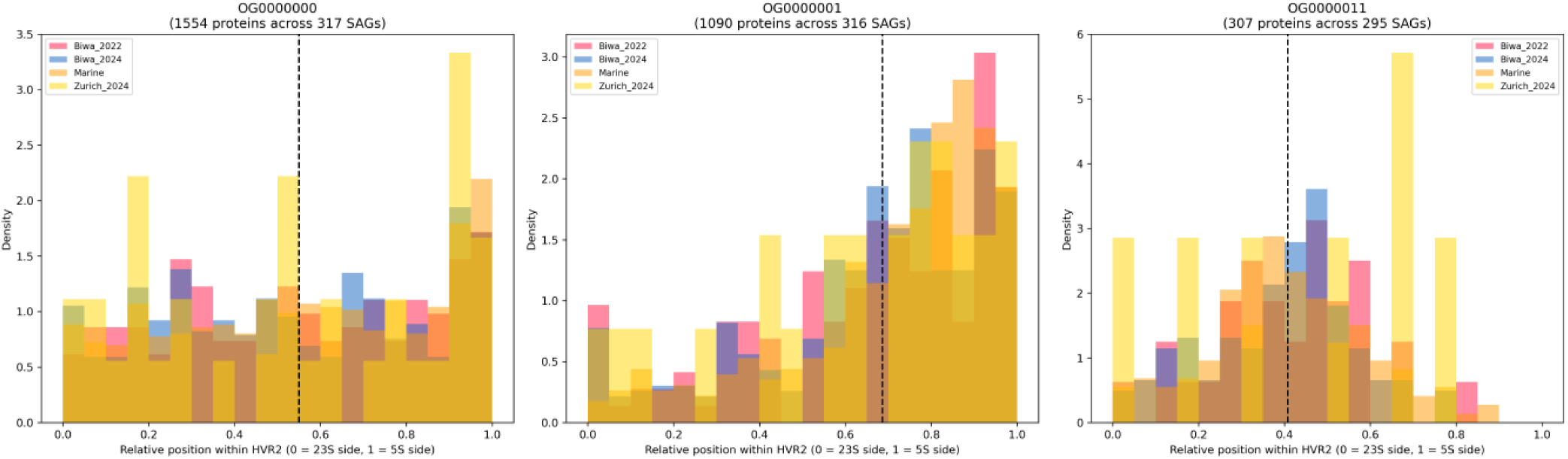
Relative genomic positions of the three core gene families (OG0000000, OG0000001, OG0000011) within HVR2 sequences. The x-axis represents the normalised position along the HVR2 length, and the y-axis represents the density of individual HVR2 sequences that contain these gene families at said position. Gene positions differ based on the family, with OG0000001 skewed towards the 5S end (mean 0.69), whereas OG0000011 and OG0000000 are distributed across the HVR2 (means of 0.41 and 0.55 respectively).

**Figure S5.**
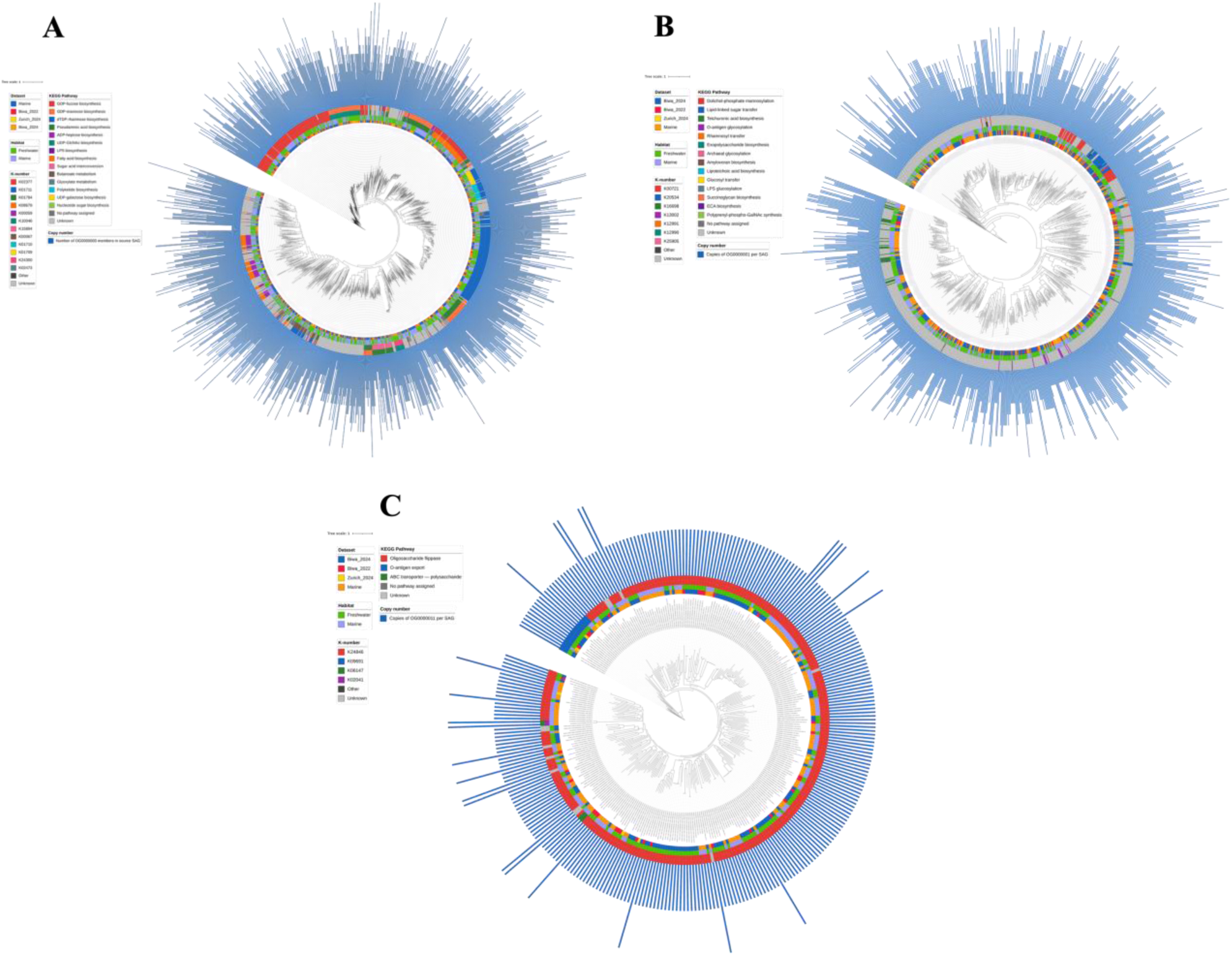
Maximum likelihood phylogenetic trees of the three core gene families (OG0000000, OG0000001, OG0000011) across marine and freshwater HVR2 sequences. (a, b, **c)** Trees were constructed using IQ-TREE 3, with sequences aligned by MAFFT. Marine and freshwater sequences are intermixed throughout each tree rather than forming habitat-specific clades.

**Figure S6.**
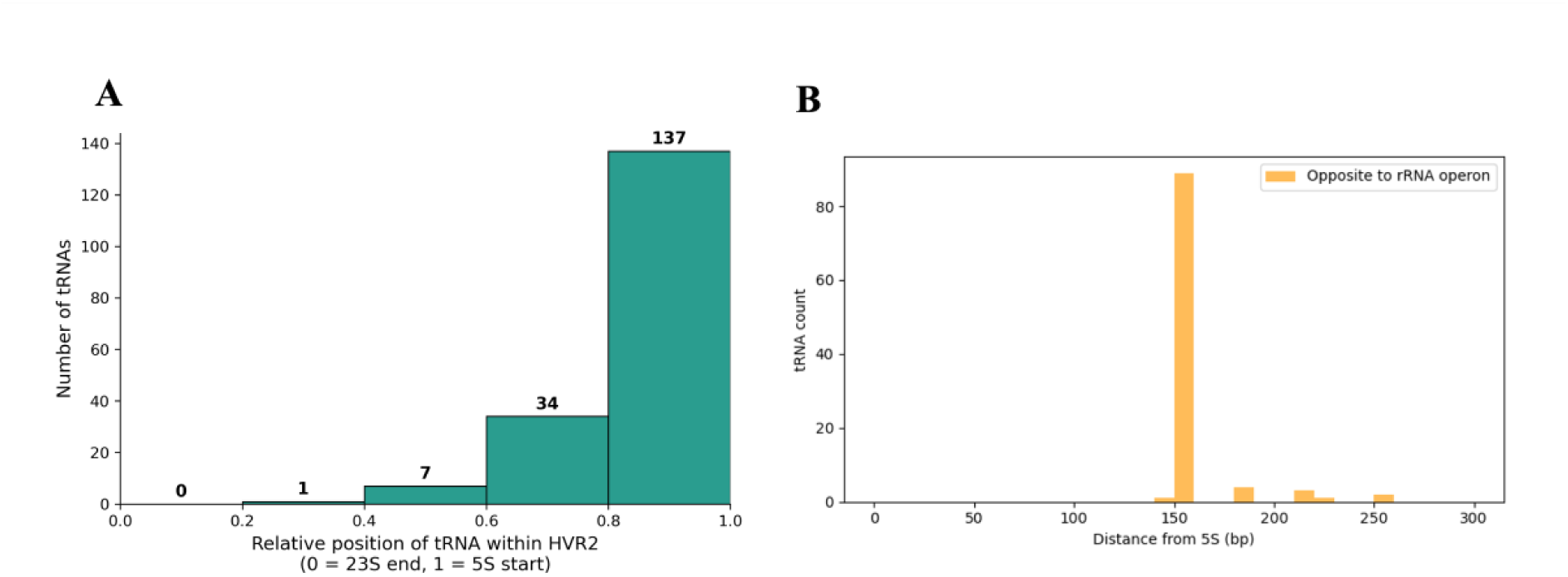
tRNA distribution in *Fontibacterium* HVR2. **(a)** Distribution of tRNA gene positions within freshwater HVR2 sequences relative to the flanking rRNA gene boundaries. The x-axis represents the distance from the nearest rRNA boundary (23S or 5S), and the y-axis represents the number of tRNA genes. A total of 179 initiator methionine tRNA genes were identified across all 176 freshwater HVR2 sequences using tRNAscan-SE with the bacterial model. The majority of tRNA genes (n = 141, 79%) were located within 10 kb of an rRNA boundary, and for almost all genes (177 of 179), the nearest boundary was the 5S rRNA gene. **(b)** Close-up distribution of the 100 tRNA genes located within 300 bp of the flanking 5S gene boundary in freshwater HVR2 sequences, where most were concentrated around the 150 bp locus.

